# Enhanced Kidney Targeting and Distribution of Tubuloids During Normothermic Perfusion

**DOI:** 10.1101/2025.04.10.648123

**Authors:** Enrique Montagud-Marrahi, Adriana Rodriguez-Gonzalo, Rubén López-Aladid, Yosu Luque, Ruben Rabadan Ros, Elena Cuadrado-Payan, Elisenda Bañón-Maneus, Jordi Rovira, Marta Lazo-Rodríguez, Oriol Aguilà, Carolt Arana, Ainhoa García-Busquets, Natalia Hierro, Thomas Prudhomme, Mireia Musquera, Yun Xia, Fritz Diekmann, Josep M. Campistol, Maria José Ramírez-Bajo

## Abstract

Tubuloids have become a promising tool for kidney disease model and regeneration, although their ability for integration and regeneration in vivo is not well documented. Here we establish, characterize and compare human tubuloids using two optimized protocols, which added a prior isolation of tubular cells (Crude tubuloids) and proximal tubular cells (F4 tubuloids). Secondly, using this protocol, healthy rat-derived tubuloids were established. Finally, we compared two strategies to deliver GFP tubuloids in a kidney host: 1) subcapsular/intracortical injection and 2) tubuloid infusion during normothermic preservation in a rat transplantation model and a human discarded kidney. F4 tubuloids achieved a more differentiation state compared to Crude tubuloids. When analyzing tubuloid delivery to the kidney, normothermic perfusion was more efficient compared to in vivo injection, observing fully-developed tubules in the host parenchyma after 1 week and 1 month. Normothermic perfusion also successfully delivered kidney tubuloids to a discarded human kidney for transplantation. These results suggest that tubuloids administered during normothermic perfusion represent a potential strategy to enhance the translatability of kidney regenerative therapies to the clinical practice to condition kidney grafts and to treat kidney diseases.

## 1. INTRODUCTION

Among the different models of kidney organoids proposed to date, tubuloids have been recently developed as a new model of kidney organoids. Kidney tubuloids are constituted by a more mature cellular structure of kidney tubular cells that mimic kidney tubules in a three-dimensional distribution and with high precision^1–3^. An important advantage of these constructs is that they can be obtained from adult kidney tissue and they are associated with less technical complexity compared to kidney organoids, since they are developed *in vitro* from cells extracted from kidney tissue (biopsies or nephrectomies) or directly from urine^2^.

In addition to in vitro disease modelling, tubuloids offer a potential strategy for renal regenerative medicine, although studies with their application *in vivo* have not yet been performed. Previous experiences have shown successful integration of kidney organoids in vivo by different techniques, although most of them only achieve a local integration and are difficult to translate to clinical practice^4,5^. Therefore, recent experiences have focused on new approaches to efficiently reach the target organ with the desired treatment. Among them, ex vivo normothermic perfusion (NMP) of solid organs has arisen as one of the most promising platforms for organ-specific treatment delivery^6–10^. This system creates a platform for administering and delivering regenerative therapies to the target organ in a controlled and efficient manner^6^.

Here, we describe two adapted protocols for human and rat tubuloid obtention performing a previous purification of tubular cells to explore tubuloid infusion during kidney NMP as a strategy for efficiently delivering tubuloids to the kidney that can be translated into clinical practice.

## 2. METHODS

### 2.1. Human tubuloid culture

Human kidney tissue for tubuloid culture was collected after nephrectomy from areas of healthy parenchyma from patients with a kidney neoplasm but a preserved kidney function. The study was approved by the Research Ethics Committee of our centre, and signed consent forms are available upon request.

To increase tubular cell purity from the whole kidney sample we include a tubular cell enrichment step as previously described by Vinay et al.^11^ The digested tissue was sequentially sieved through a 320 µm and 150 µm sieves and then resuspended in Matrigel^®^ (Corning) and plated to obtain tubuloids (protocol A, Crude tubuloids). We performed a second protocol (protocol B, F4 tubuloids) in which the tissue further underwent a second separation process by centrifugation with 45% Percoll (Figure 1A). See Supporting Methods, Human tubuloid culture.

**Figure 1.**
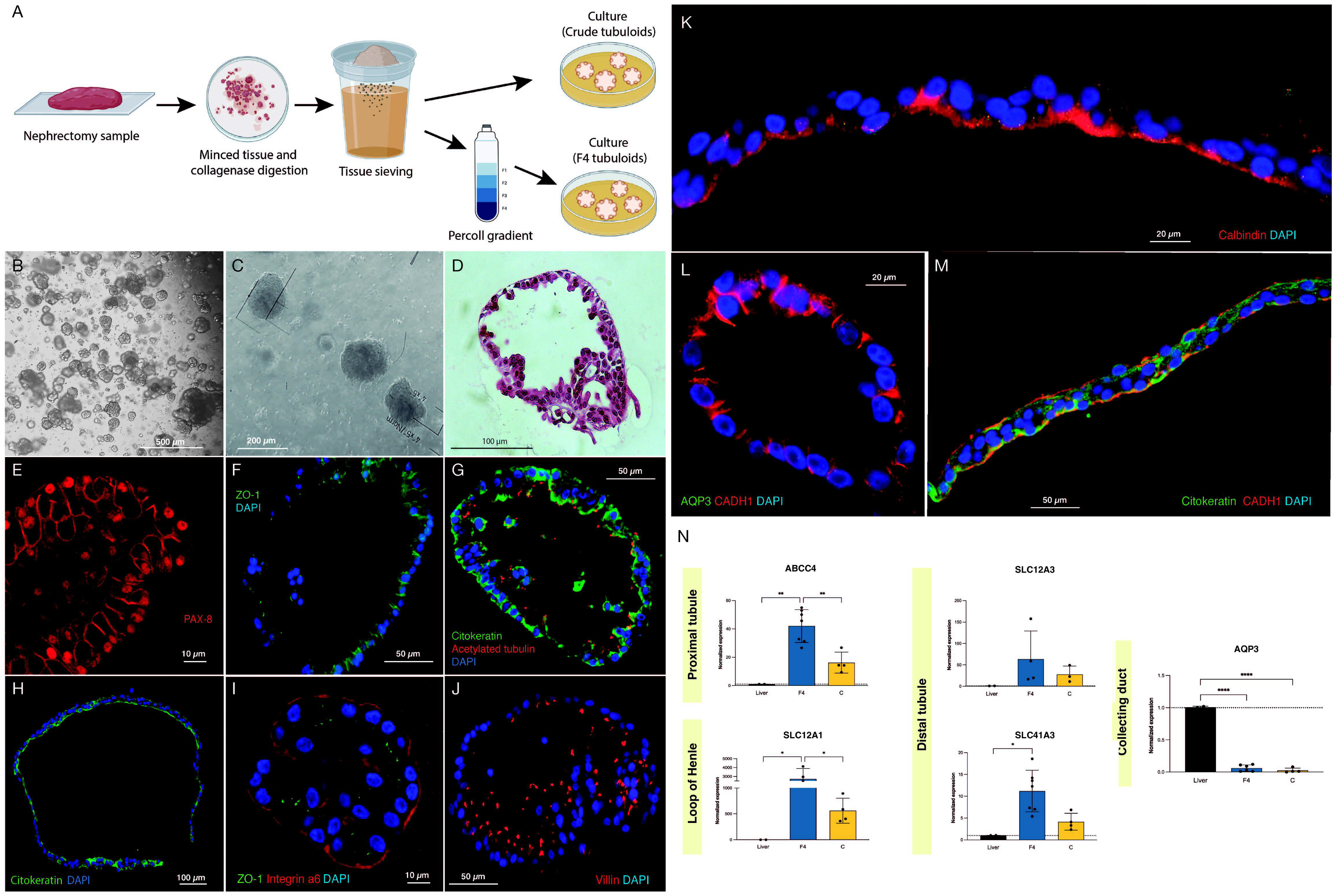
Crude and F4 kidney human tubuloids show typical tubular polarity and specific markers of different nephron segments. A, Methods for Crude and F4 obtention. B-C, representative images from optical microscopy of human F4 (B) and Crude tubuloids (C) showing a spheroid morphology (obtained from n = 7 independent kidney tissue donors). D, Paraffin sections with HE-staining of a fully-developed F4 human kidney tubuloid with a single cell layer and a spheroid shape (n = 7 independent kidney tissue donors). E, PAX-8 immunofluorescence in a Crude human tubuloid. F, immunofluorescence for ZO-1 in a F4 tubuloid. G, immunofluorescence for pan-cytokeratin and acetylated tubulin in a F4 tubuloid. H, immunofluorescence for pan-cytokeratin in a Crude tubuloid. I, immunofluorescence for Integrin a6 and ZO-1 in an F4 tubuloid. J, immunofluorescence of Villin in an F4 human tubuloid. K, Crude tubuloid showing a positive staining for Calbindin. L (Crude tubuloid) and M (F4 tubuloid), immunofluorescence for AQP3, CADH1 and cytokeratin. Tubuloids were obtained from n = 7 independent kidney tissue donors, and each condition was repeated in duplicate. N, Expression of tubular markers in human tubuloids. Black box: liver tissue; Blue box: F4 tubuloids; Yellow box: Crude tubuloids. *P < 0.05, **P < 0.001, ****P < 0.0001. Data are represented as mean ± SEM. For qPCR analysis, n = 7 independent kidney tissue donors for F4 tubuloids and n = 4 independent kidney tissue donors for Crude tubuloids (matched with F4 tubuloids’ donors). Triplicates were performed for each sample and condition.

### 2.2. Rat tubuloid culture

Rat kidney tissue for rat tubuloid culture was collected from healthy 3 months-old Lewis rats (LEW/Han®Hsd, Envigo^®^). Tubuloid isolation and culture were performed following the same methods that those described for human tubuloids. The study was approved by and conducted according to the guidelines of the local animal ethics committee (Comité Ètic d’Experimentació Animal, CEEA, Decret 214/97, Catalonia, Spain). See Supporting Methods, Rat tubuloid culture.

### 2.3. GFP tubuloids development

For tracking tubuloids in delivery studies, GFP-expressing rat tubuloids were obtained from a GFP rat strain (CAG-GFP rat line, genOway) with a constitutive expression of GFP. The method performed for isolation and culture of GFP rat tubuloids from this strain was similar to that previously detailed. Both protocols were performed (F4 and Crude). GFP fluorescence was confirmed immediately after plating with a fluorescence microscope.

### 2.4. Subcapsular and intracortical tubuloid injection in vivo

GFP-expressing rat tubuloids were harvested and resuspended in pure Matrigel® without previous disruption. A Lewis rat was selected as a host ((LEW, RT1-A^1^, Envigo). The animal was anesthetized with inhaled isoflurane and an abdominal middle incision was performed to expose the abdominal organs. Around 1·10^6^ cells were injected in the left kidney: two injections were performed (one subcapsular in the upper kidney pole and the other intracortical in the lower pole) with a 25G needle. Rats were euthanized 1 day after the procedure and both native kidneys were collected for analysis.

### 2.5. Tubuloid infusion during rat kidney normothermic perfusion

Rat donor organ procurement was performed under anesthesia with isoflurane (IsoVet®, Braun) as previously described^12^. Rat was performed following the protocol published elsewhere^13^.

GFP-expressing rat tubuloids were harvested and resuspended in pure Matrigel® with previous mechanical disaggregation and then were administered in the perfusate through the arterial line. Kidney grafts were maintained under NMP for 1 hour. For each donor, one kidney graft was perfused and treated with tubuloids, while the contralateral one was perfused but no tubuloids were administered. We tested two doses of tubuloids: 1·10^6^ cels/g (high dose, n = 4 kidneys) and 3·10^5^ cels/g (low dose, n = 3 kidneys). See Supporting Methods, Tubuloid infusion during rat kidney normothermic perfusion.

### 2.6. Kidney transplantation in rats

Inbred 3-months Lewis rats were used as recipients for kidney grafts preserved in NMP. Since the CAG-eGFP rat strain had a Sprague-Dawley background, a short dose of tacrolimus was established to avoid a potential rejection response against GFP tubuloids. Thus, 0.5 mg/kg of tacrolimus (Modigraf®) was administered intramuscularly to the Lewis recipient during four consecutive days (−1, 0, +1 and +2 concerning transplantation – day 0)^14^. Kidney transplants were performed as previously described by our group^12^. See Supporting Methods, Kidney transplantation in rats.

### 2.7. Tubuloid infusion during human kidney normothermic perfusion

Human kidneys were obtained from a patient who was declared as a potential donor and the organs were evaluated according to our center policy. Only after the decision to be discarded according to clinical criteria, kidneys were accepted for research purposes. The study protocol was reviewed and approved by the Research Ethics Committee of our centre (Comité de Ética de la Investigación con medicamentos, CEIm). Written informed consent to participate in this study was provided by the patient’s relatives.

Normothermic Perfusion was performed using the ARK Kidney device from Ebers Medical® as previously described by our group^10^. Both kidneys from the same donor were simultaneously perfused, and GFP-rat tubuloids (1·10^6^ cels/g) were administered in one of them through the arterial line once the organ was stable. Both kidneys were maintained for 6 hours.

For a more detailed information regarding Methods, see Supporting Information.

### 2.8. Statistical Analysis

Statistical Analysis was performed using Student T-test or ANOVA test accordingly. Data was represented in graphs using GraphPad Prism version 9.0.0 for Mac (GraphPad Software, San Diego, California USA). A P < 0.05 was considered significant.

## 3. RESULTS

### 3.1. Crude and F4 kidney human tubuloids show typical tubular polarity and specific markers of different nephron segments

Human tubuloids typically developed about 7 days after plating. Optical microscopy (Figure 1B and C) and Hematoxilin and Eosin (Figure 1D) showed that fully developed tubuloids acquired a typical morphology of a polarized epithelium with a single cell layer and a spheroid shape, usually observed after 3-4 passages. No significative differences in culture and development were observed among Crude and F4 tubuloids.

To assess the epithelial tubular nature of the obtained tubuloids, a specific immunostaining against PAX-8 was performed, showing a sustained and general expression of this marker in tubuloid cells, mostly in nuclei (Figure 1E). After assessing the tubular nature, we further analyzed the presence of a typical tubular polarization of the epithelium: ZO-1 (a tight junction tubular protein) was predominantly situated on the lateral and apical sides (Figure 1F). Polarization was also evidenced by combining immunofluorescence of pan-cytokeratin (basolateral) and acetylated tubulin (apical) (Figure 1 G-H). Integrin α6 (which maintains the integrity of the kidney tubular epithelium) was mostly expressed in the basolateral side of the tubuloids’ cells (Figure 1I).

Immunostaining of Villin 1 (apical) identified proximal tubular cells (Figure 1J). Distal cells were objectified by positive immunostaining for Calbindin-1 (an intracellular calcium-binding protein that is localized in the distal tubule cells) on the apical side (Figure 1K). E-cadherin (CADH1), a basolateral marker for the adherens junctions, is typically present in distal and collecting duct cells. Nevertheless, in collecting duct cells, CADH1 is usually expressed with AQP3 (a channel expressed only in collecting duct cells). We identified the presence of CADH1-positive tubuloids, although no concomitant expression of AQP3 was evidenced, thus suggesting a distal nature of these tubular cells (Figure 1 L-M). Remarkably, those tubuloids expressing a specific nephron segment marker did not express another specific marker for other segments, suggesting a predominant tubular cell type constituting the tubuloid. These findings were observed in either Crude or F4 tubuloids.

Proximal gene ABCC4 expression was higher when compared to the liver, as well as for SLC12A1 (loop of Henle), SLC12A3 and SLC41A3 (distal tubule). AQP3 (collecting duct) was not significantly expressed in human tubuloids. F4 tubuloids exhibited a higher expression of every tubular marker compared to Crude tubuloids (Figure 1N).

### 3.2. Expanding F4 kidney human tubuloids are constituted by cells with a polarized proximal and distal tubular phenotype, compared to Crude tubuloids

To further investigate the different cell types present in the developed human tubuloids through our two methods, we performed a single-cell RNA (scRNA) sequencing of Crude Tubuloids (n = 4, 49442 cells, Passage 3 and 4) and F4 Tubuloids (n = 3, 29998 cells, Passage 3 and 4). The clustering analysis resulted in the identification of 2 distinct clusters in both Crude and F4 tubuloids (Cluster 0 and Cluster 1)(Supporting Table S1).

For Crude tubuloids both clusters exhibited an overlapping of tubular cell markers expression: Cluster 1 displayed a predominant proximal tubular cell signature (marked by expression of NHS, FHIT and PTPRM^15,16^), also a minor expression of distal tubular markers was identified (represented by CACNA2D3^15,16^). This pattern was lacking in Cluster 0, although a reduced expression of almost every marker tested was also present. This minor expression of segment-specific genes may suggest a population with a more progenitor status. In both clusters, a minor population of cells with a collecting duct signature (marked by NALF1 and RPA1^15,17^) were identified (Figures 2A-D).

**Figure 2.**
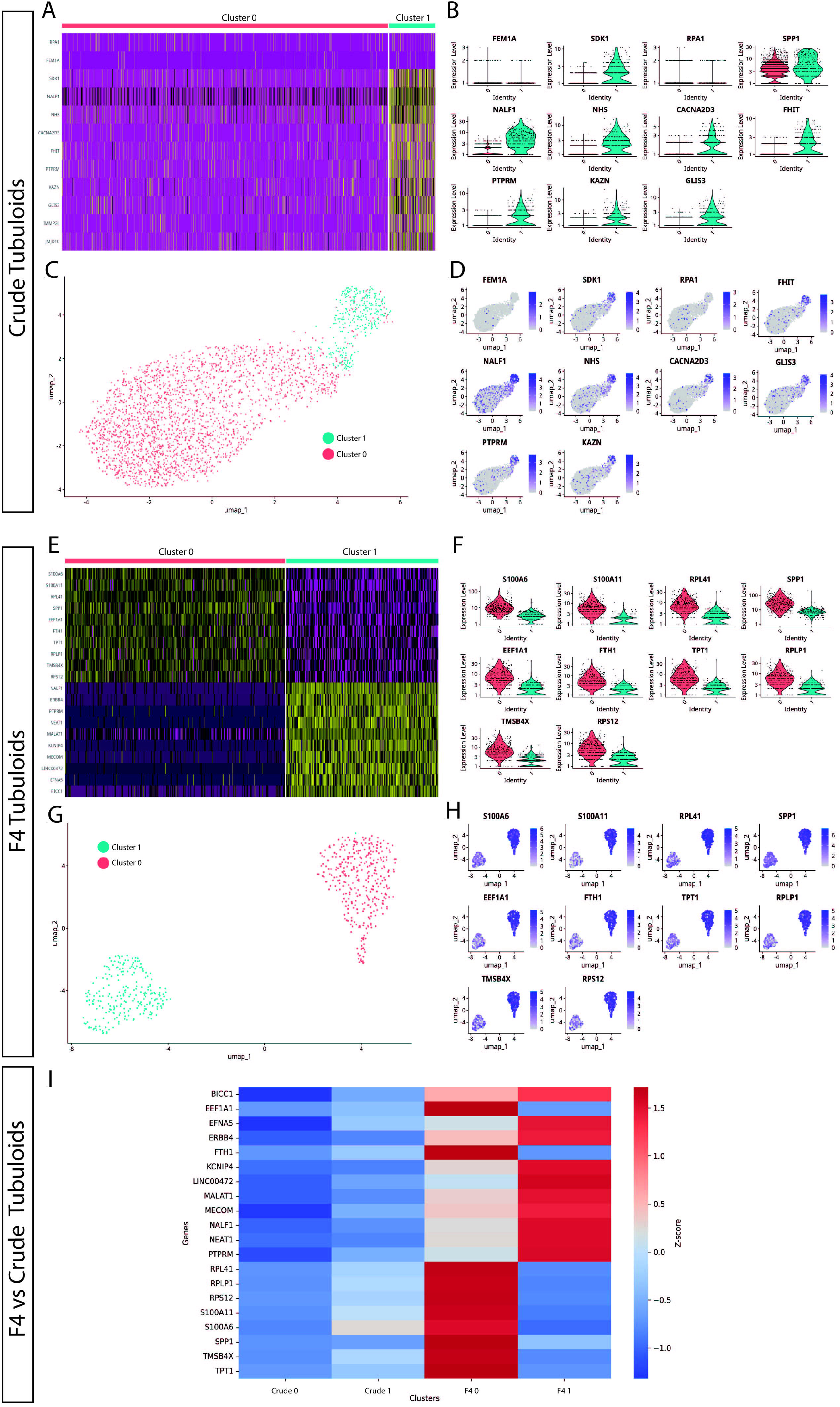
Expanding F4 kidney human tubuloids are constituted by cells with a polarized proximal and distal tubular phenotype, compared to Crude tubuloids. A, Heatmap showing gene expression pattern in two cell clusters for Crude tubuloids. B, Differentially expressed genes between the identified clusters in Crude tubuloids. C, Uniform manifold approximation and projection (UMAP) plot and clustering of single-cell RNA sequencing data of Crude tubuloids (n = 4, 49442 cells, Passage 3 and 4). D, Feature plot for the differentially expressed genes for Crude tubuloids. E, Heatmap showing gene expression pattern in two cell clusters for F4 tubuloids. F, Differentially expressed genes between the identified clusters in F4 tubuloids. G, UMAP plot and clustering of single-cell RNA sequencing data of F4 tubuloids (n = 3, 29998 cells, Passage 3 and 4). H, Feature plot for the differentially expressed genes for F4 tubuloids. I, Heatmap of normalized Z-scores for expressing proximal and distal tubular markers across Crude and F4 tubuloids. See also Supporting Table 1.

Two cell clusters were also identified in F4 tubuloids with significantly different expression signatures: Cluster 0 displayed a predominant proximal tubular cell signature (marked by FTH, SPP1, TPT1 and RPLP1^15,16^), whereas Cluster 1 mostly depicted a distal tubular cell signature (marked especially by KCNIP4 and NEAT1^15,16^)(Figure 2 E-H). Noticeably, both clusters showed a clear separation of gene expression for specific-segment genes, suggesting a higher differentiation grade than Crude tubuloids. Cluster 0 also showed a marked expression of S100A6 (involved in calcium management), which has been related to tubuloid-derived progenitors from the thick ascending limb^3^. Furthermore, F4 tubuloids depicted a high expression of genes involved in protein synthesis and cell proliferation (RPS12, MECOM, BICC1^15,16^), thus indicating a high proliferation status of F4 tubuloids.

When gene expression for tubular markers of both culture protocols was confronted through a Z-score normalization, we observed that Crude tubuloids exhibited negative Z-scores for the majority of analyzed genes indicating a low relative expression to the overall mean across conditions. This was especially marked for Crude Cluster 0, while Cluster 1 depicted higher expression levels. In contrast, F4 tubuloids were associated with predominantly positive Z-scores, suggesting a higher level of expression for the analyzed genes. Furthermore, a polarized expression for proximal (F4 Cluster 0) or distal (F4 Cluster 1) tubular markers was observed. This differential pattern indicates that the F4 culture method may provide an environment more conducive to gene expression changes associated with differentiation (Figure 2I).

### 3.3. Tubuloids can be obtained from adult rat kidney tissue and represent a new in vitro platform for studying kidney diseases

Rat tubuloids developed after 10-14 days and acquired a uniform spherical morphology, which was preserved for up to 17 passages (Figure 3A and B). Immunofluorescence assessment evidenced a tubular epithelial polarity, similar to the human ones (Figure 3C and D).

**Figure 3.**
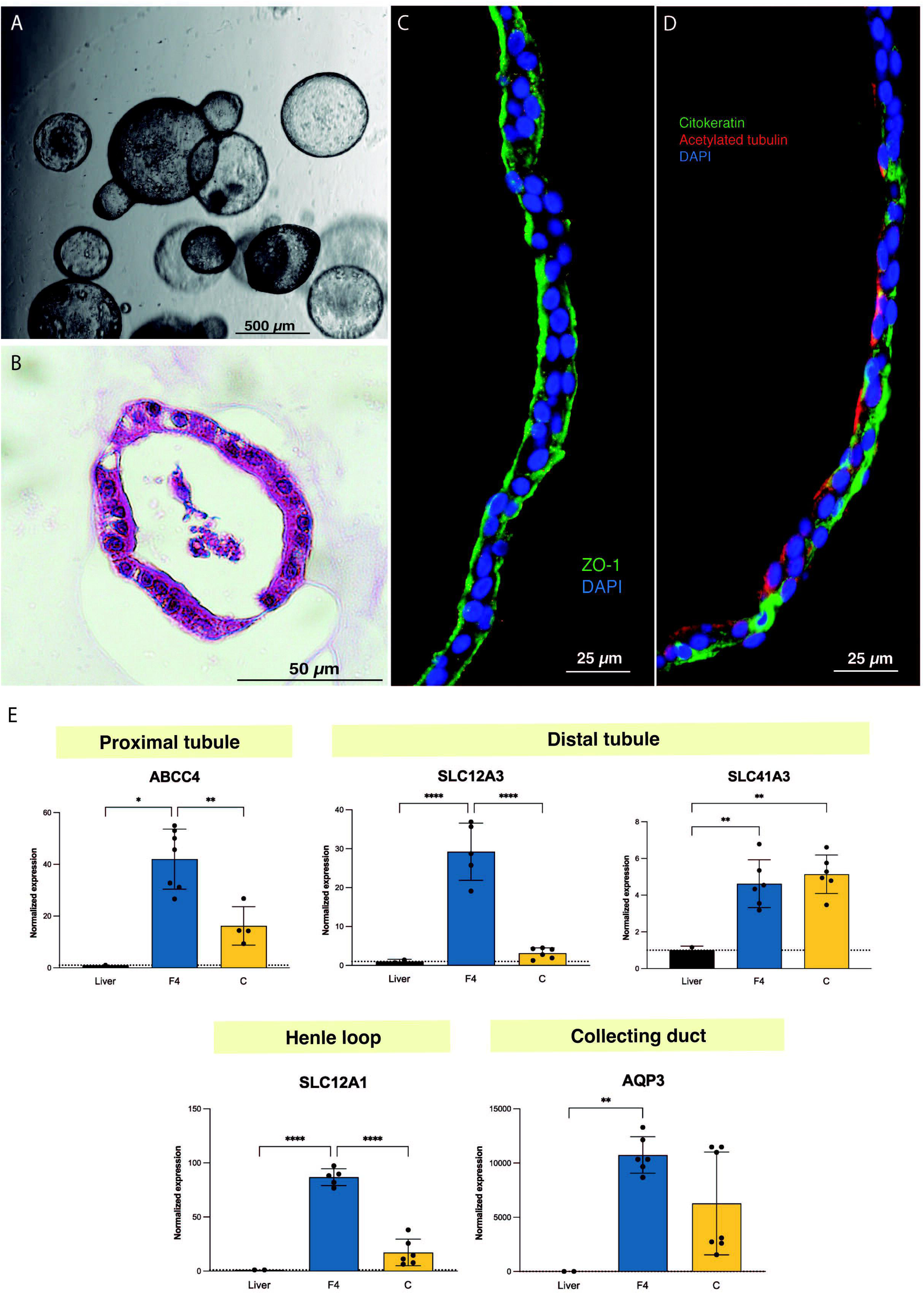
Tubuloids can be obtained from adult rat kidney tissue. A, Optical microscopy image of rat tubuloids showing a uniform spherical morphology. B, Paraffin sections with HE staining of rat tubuloids. C and D, Confocal images of rat tubuloids showing an epithelial polarization as human tubuloids for ZO-1 (C), cytokeratin and acetylated tubulin (D). E, Expression analysis of rat tubuloids for different tubular genes. Black box: liver tissue; Blue box: F4 rat tubuloids. Yellow Box: Crude rat tubuloids. *P < 0.05, **P < 0.001, ****P < 0.0001. Data are represented as mean ± SEM. Rat tubuloids were obtained from n = 8 independent Lewis rats, seeded per duplicate to obtain matched F4 and Crude tubuloids. For qPCR analysis, liver tissue was obtained from 2 independent Lewis rats. Each sample was analysed per triplicate for each condition.

Gene expression analysis for segment-specific markers evidenced the expression of proximal, Loop of Henle and distal tubule markers, especially for F4 rat tubuloids. In contrast to human tubuloids, rat ones expressed the collecting duct marker AQP3 (Figure 3E).

### 3.4. Kidney ex vivo normothermic perfusion provides an efficient strategy for tubuloid delivery to kidney parenchyma

The epifluorescence assessment of the in vivo injected kidney with the IVIS® Lumina device evidenced a positive GFP signal in the injected kidney compared to the contralateral one (without treatment). This signal was localized in the kidney surface where the injections were performed (Figure 4A-C). No evident signal was detected in the internal part of the kidney. The immunofluorescence analysis showed the presence of GFP-positive structures of 150-200 μM of diameter localized in the injection area (Figure 4D).

**Figure 4.**
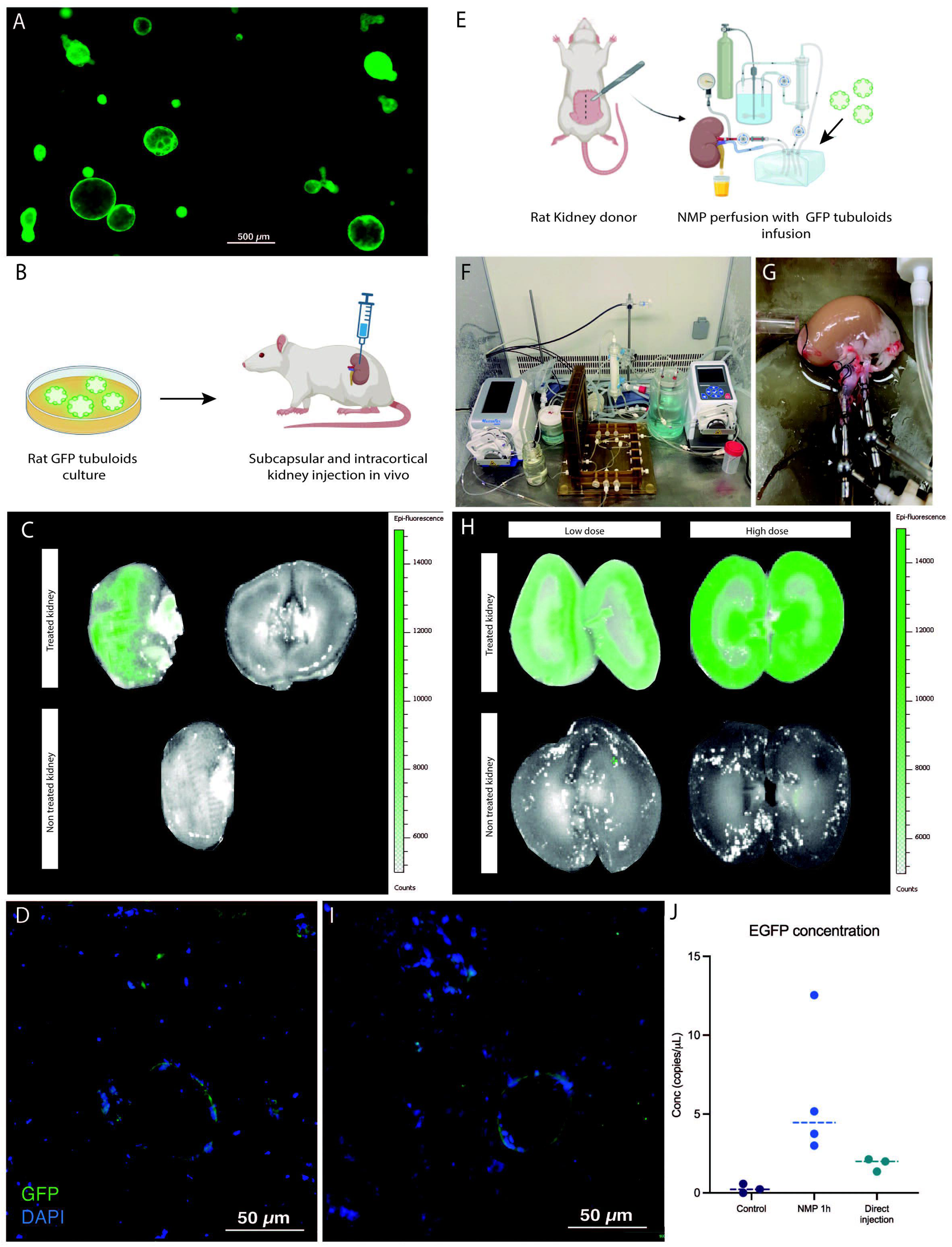
Kidney ex vivo normothermic perfusion provides an efficient strategy for tubuloid delivery to kidney parenchyma compared to in vivo administration. A, Fluorescence Optical Microscopy image from rat tubuloids obtained from CAG-eGFP rat kidney tissue (obtained from n = 5 independent kidney rat donors). B, Scheme of in vivo injection of GFP tubuloids in a rat kidney host. C, Epifluorescence analysis of GFP signal in the injected kidney with GFP tubuloids (n = 3 injected rats). Image is showing the outer and internal parenchyma of one representative case of an in vivo injected kidney, as well as a control one (no injection). D, Representative image of a GFP tubuloid observed in the kidney parenchyma after subcapsular injection (n = 3 injected rats, each immunofluorescence analysis was performed in duplicate). E, Scheme of the GFP tubuloid injection during kidney normothermic preservation. F-G, Detail of the normothermic system for kidney rat normothermic preservation. H, IVIs images for GFP fluorescence of perfused and treated kidneys with high (n = 4 kidneys from independent Lewis donors) and low doses (n = 3 kidneys from independent Lewis donors) of tubuloids and perfused, non-treated kidney grafts (n = 3 kidneys from independent Lewis donors). I, Representative immunofluorescence image for GFP showing a GFP tubuloid in the kidney cortical after normothermic perfusion and tubuloid infusion (high dose, n = 4 kidneys from independent donors, each condition performed in duplicate). J, dPCR eGFP copies determination for kidneys perfused with GFP tubuloids and those in which direct injection was performed. Data are represented as individual values and median. Control, n = 3 perfused, non-treated kidneys. NMP 1h, n = 4 perfused and treated kidneys, high dose. Direct injection, n = 3 injected rats. For dPCR, each sample was analyzed per duplicate for eGFP expression. Epifluorescence and immunofluorescence images were acquired under the same acquisition and detection settings, and a protocol to mitigate kidney autofluorescence was performed. GFP signal was corrected for background.

The second strategy consisted of infusing GFP tubuloids in the perfusate during ex vivo normothermic perfusion of the kidney graft (Figure 4E-G). We tested two cell doses to assess a potential correlation with the fluorescent signal observed on the IVIs analysis (high dose: 1·10^6^, and low dose: 3·10^5^ cells/g). Tubuloid infusion was usually followed by a temporary increased vascular resistance (Supporting Figure 2). IVIs study evidenced a diffuse GFP signal mostly in the kidney cortex, with no significant signal in the perfused, non-treated kidney (Figure 4H). The registered fluorescence was higher and acquired a more diffuse distribution than the in vivo tubuloid injection (Figure 4C and H). When testing a lower dose of tubuloids, a similar pattern, although with a decreased signal, was identified in the treated kidney graft (Figure 4H). The immunofluorescence analysis showed the presence of GFP-positive structures of 30-50 μM diameter in the cortical kidney parenchyma (Figure 4I).

Kidney grafts treated with tubuloids during NMP presented the highest number of EGFP copies, with a concentration that ranged from 3 to 12 copies/μL. In vivo injected kidney grafts showed fewer EGFP copies (ranging from 2 to 3 copies/μL), although higher than the control (Figure 4J).

### 3.5. Tubuloids delivered to the kidney during NMP integrate into the host kidney parenchyma and form new tubular structures

At 1 week and 1 month, the kidney graft presented a positive GFP signal, mostly in the cortex (Figure S1, S2 and Figure 5A and 5B). Noticeably, GFP fluorescence decreased compared to kidney graft just after NMP, even more after 1 month. No signal was detected either in the contralateral native kidney or the liver at the analyzed timepoints (Figure 5B). DdPCR analysis for GFP copies evidenced a higher concentration in the treated grafts compared to controls that persisted at 1 month (Figure 5C). Immunofluorescence for GFP in the kidney graft showed the presence of GFP-expressing cells in the tubular compartment (Figure 5D). Remarkably, those cells constituted new fully-developed tubules in the host kidney parenchyma but were also found integrating the pre-existing epithelium of host tubules. Those newly developed tubules showed a positive expression of Villin, while expression of Calbindin-1 was not observed, thus suggesting a proximal phenotype with no formation of distal tubules (Figure 5D).

**Figure 5.**
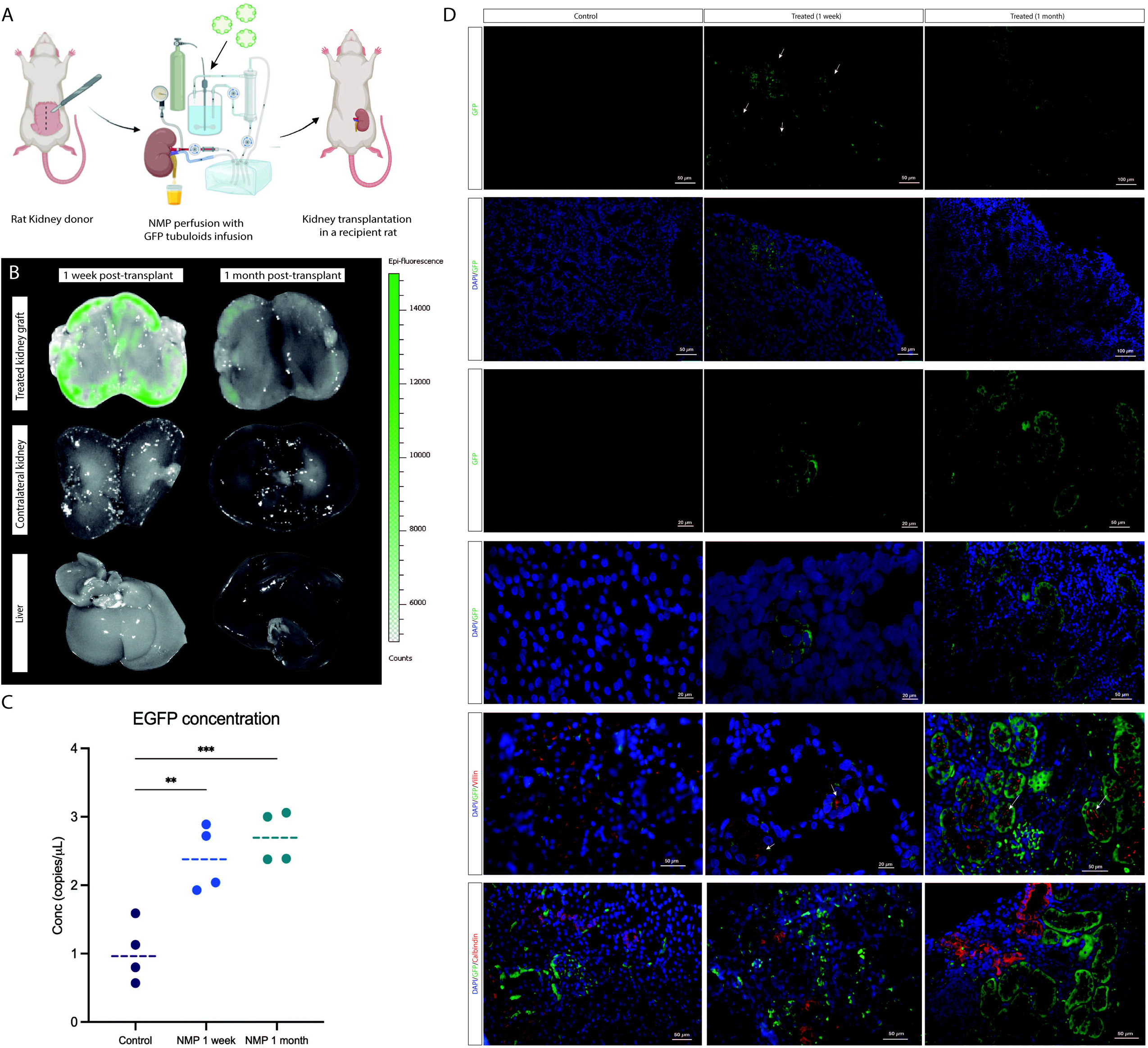
Tubuloids delivered to the kidney during NMP integrate into the host kidney parenchyma and form new tubular structures. A, Scheme of the GFP tubuloid injection during human kidney normothermic preservation and transplantation. B, IVIs images for GFP fluorescence of the treated kidney graft, native kidney and liver. C, dPCR eGFP copy number determination for kidneys perfused with and without GFP tubuloids. D, Immunofluorescence images for GFP, Villin and Calbindin-1 of the control and treated kidneys showing GFP and Villin-expressing cells (white arrows) constituting new tubuloids but also integrated in the host tubular epithelium after normothermic perfusion and tubuloid transplantation. Data are represented as individual values and median. *P < 0.05. ** P < 0.01. *** P < 0.001. N = 4 transplanted rats per group (control, 1 week and 1 month). For immunofluorescence and dPCR studies, each sample was performed in duplicate. Epifluorescence and immunofluorescence images were acquired under the same acquisition and detection settings, and a protocol to mitigate kidney autofluorescence was performed. GFP signal was corrected for background.

### 3.6. Ex vivo normothermic perfusion efficiently delivers tubuloids in a human kidney graft

The kidney donor was a 56-year-old man without any relevant medical history who experienced a cardiac arrest with no recovery after 90 minutes of basic and advanced vital support. Once identified, organ retrieval was immediately started as an Uncontrolled Donor after Cardiac Death (uDCD). Serum creatinine at donation was 1.5 mg/dL, although bad perfusion of both kidneys was observed in the bench surgery, thus being discarded for transplantation.

Both kidneys were simultaneously cannulated and connected to the Ebers® NMP perfusion device (Figure 6A and B). After stabilization, GFP tubuloids were mechanically disaggregated until 50 μm structures and then administrated in one of the kidneys through the arterial cannula at a dose of 1·10^6^ cells/g. Kidneys were perfused for 6 hours. Hemodynamics during perfusion are depicted in Supporting Figure S3.

**Figure 6.**
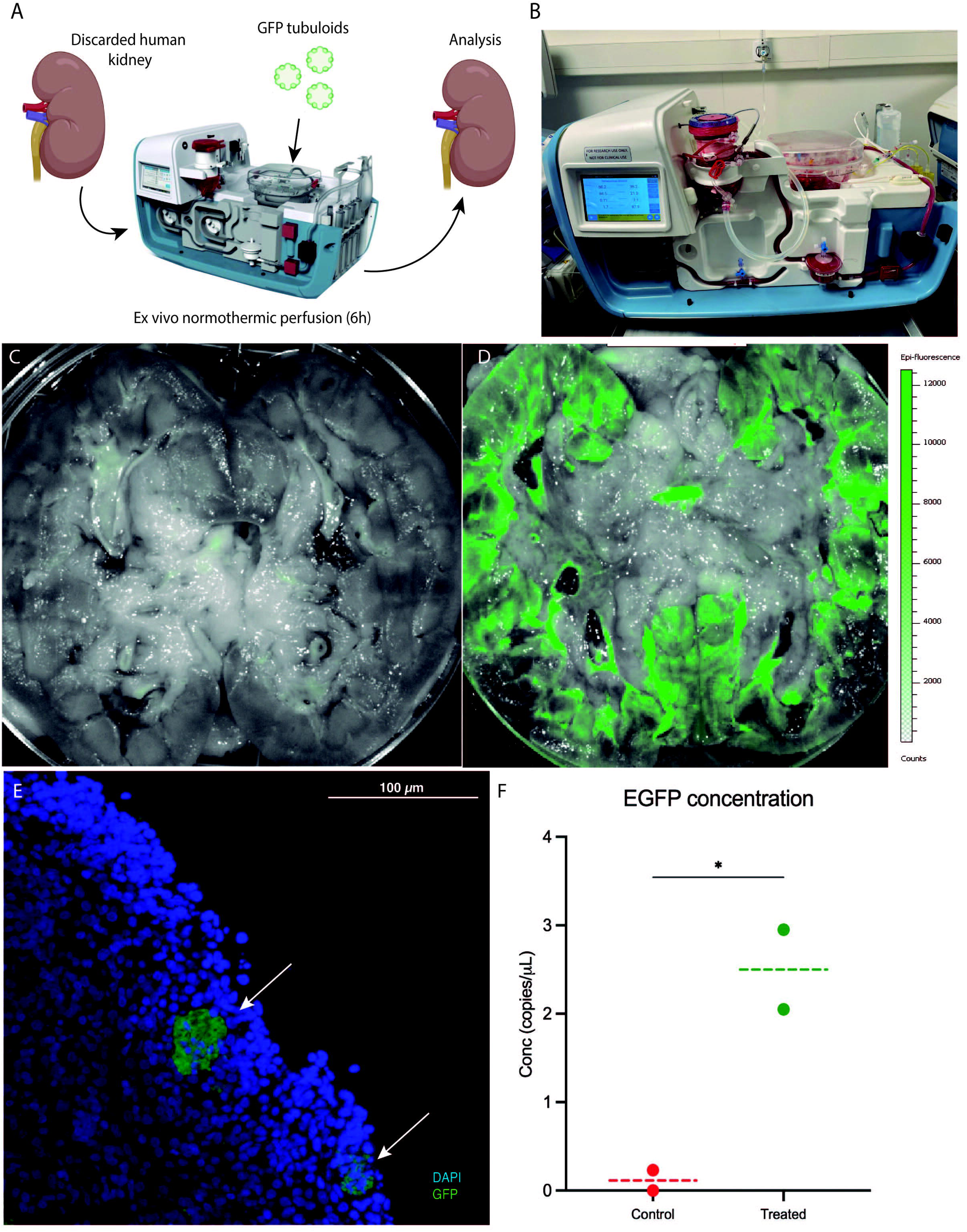
Ex vivo normothermic perfusion efficiently delivers tubuloids in human kidney grafts. A, Scheme of the GFP tubuloid injection during human kidney normothermic preservation. B, Detail of the normothermic system for human kidney normothermic preservation. C and D, IVIs images for GFP fluorescence of perfused kidneys without (C) and with GFP tubuloids (D). E, Representative immunofluorescence image for GFP showing GFP tubuloids (white arrows) in the human kidney after normothermic perfusion and tubuloid infusion. F, dPCR eGFP copy number determination for treated and non-treated human kidneys with GFP tubuloids during NMP. Each dot represents one random sample from the same kidney parenchyma. Line represents the median. N = 1 kidney per group (control and treated). Each sample was processed in duplicate for dPCR and immunofluorescence analysis. Epifluorescence and immunofluorescence images were acquired under the same acquisition and detection settings, and a protocol to mitigate kidney autofluorescence was performed. GFP signal was corrected for background.

After 6 hours, IVIs assessment showed a positive and diffuse GFP fluorescence sign in the perfused, treated kidney, mostly located in the kidney cortex, with no significant signal in the perfused, non-treated kidney (Figure 5C and D). Immunofluorescence assessment evidenced the presence of 20 to 50 μm GFP-positive cell aggregates in the treated kidney cortical graft. GFP copy quantification in the kidney tissue showed a higher concentration in the treated graft compared to the control (2.5 vs 0.1 copies/μL, respectively). Expression of kidney injury (KIM-1), apoptosis (CASP3) and inflammation markers (VEGF, TNFa) at 6 hours was lower in the treated organ, compared to the control one (Supporting Figure S4).

## 4. DISCUSSION

Even though kidney transplantation is the treatment of choice for patients with End Stage Kidney Disease (ESKD), access to a kidney graft is severely limited by the scarcity of organs available for transplantation^18^. Thus, new therapies to increase the donor pool are urgently required. In this sense, regenerative medicine encompasses a broad spectrum of strategies with the potential to overcome organ scarcity through kidney regeneration^7,19–21^. Herein, we propose two modified protocols to obtain human tubuloids from human kidney tissue by adding a tissue sieving step for isolation of tubular cells before culture. The F4 method also involved a Percoll gradient after sieving to obtain a purer tubular proximal cell population.

The resulting tubuloids from both methods expressed segment-specific markers of proximal, loop of Henle and distal tubule, but not for collecting duct. These findings were in accordance with the immunofluorescence analysis, in which no expression of AQP3 was evidenced, thus suggesting that collecting duct cells were underrepresented^2,22,23^. Remarkably, when assessing the presence of different nephron segments by immunofluorescence, each tubuloid (even those with the same origin) tended to be constituted by cells with a predominant nephron segment phenotype, a feature that has been previously described^22,24–26^.

Single-cell analysis of both culture methods showed a mixed expression of proximal, tubular and collecting duct markers, something that suggests a mixture of different cell populations. Noticeably, the absence of a specific expression of nephron markers suggests a cell population in Crude tubuloids with a more progenitor status^3^. In contrast, F4 tubuloids were composed of two cell clusters with two clearly differentiated phenotypes: one cluster was integrated by a cell population with a high expression of proximal tubular markers, while the other one expressed mostly distal tubular markers. A clear expression pattern polarization was observed in F4 tubuloids compared to Crude tubuloids, in which cell clusters overlapped. These results may be explained by the Crude tubuloids’ origin, since in this case all tubular cell types were cultured. This mixture of different tubular cells when culturing Crude tubuloids may lead to a higher number of tubular cell phenotypes with less tubuloid cell nephron-specific differentiation^3,27^.

To date, most of the studies have relied on the obtention of human tubuloids, which have received important attention^1–3^. Furthermore, the development of mice-derived tubuloids has recently been described as a new model for studying kidney disease^26,28–30^. Nevertheless, rather than mice, rats have been demonstrated to be a more reliable and advantageable model for research in kidney disease, since they have larger organs, (facilitating surgical procedures and sample collection and allowing for more precise studies), their renal physiology and pathology more closely resemble that of humans (improving the relevance of experimental findings) and rats have a slower disease progression, enabling long-term kidney function and regeneration studies. Thus, and to provide a new tool for kidney disease study, we developed rat-derived tubuloids, following the same modified protocols but using specific rat growth factors. In contrast to human tubuloids, healthy rat-derived tubuloids developed later and expressed markers of every nephron segment, even from the collecting duct. These findings may suggest a higher differentiation capacity of rat tubular cells under this specific environment and may pose this model as an available tool for further research in kidney disease^1,2,22^.

The proliferation capacity of tubuloids confers the potential ability to regenerate kidney tissue and accelerate tissue repair in kidney injury^31^, although an optimal route to be delivered and engrafted to the kidney tissue has not been established, especially when considering clinical applicability^4,5,32–36^. Recently, NMP in kidney organ transplantation has arisen as a promising platform for long-term organ preservation and treatment testing while the organ is physiologically preserved and monitored^37–39^. One of this platform’s most relevant characteristics is its fast translatability to clinical practice, since it allows testing innovative treatments in human organs and monitoring their effects in a safe environment^7,40,41^.

In our study, epifluorescence was more intense and diffuse for those grafts treated during NMP compared to those injected in vivo, in which the signal was weaker and localized in the injection area. A weaker signal was also observed when a lower dose of tubuloids was administered during NMP, suggesting a signal-cell dose relationship. Our findings were also reinforced by GFP copy quantification, since the NMP kidneys showed a copy number of up to 12- and 5-fold compared to control and injected kidneys, respectively.

In our proposal, at 1 month, GFP-expressing cells were still detected, either constituting fully-developed tubules but also side-by-side constituting the tubular epithelium among host tubular cells. Similar results were reported by Sampaziotis et al^6^ in cholangiocyte-derived organoids. Our results further evidence the engraftment capacity of tubuloids not only to integrate in a pre-existing tubular epithelium but also to generate new fully tubular structures among the host kidney parenchyma with a predominant proximal phenotype. No signal was identified in either the native kidneys or the liver after 1 month, suggesting that our approach significantly increases specific delivery to the kidney while minimizing cell leaking to non-target organs. In our scenario, cell aggregates of 20-25 μm were injected, and considering the fact that glomerular capillaries are only 6-10 μm, this may lead to blockages and severe disruption of perfusion. Nevertheless, we only observed a transitory increase in vascular resistance when tubuloids were infused, followed by a normalization of kidney hemodynamics and no vasculature blocking. We hypothesize that this observation is related to an increase in glomeruli and endothelial permeability associated with ischemia-reperfusion injury, thus allowing the transition of cellular aggregates^42–44^.

When moving to a closer clinical scenario in a discarded human kidney, we observed that, after 6 hours of perfusion, a diffuse GFP signal in the cortex was detected. Tubuloid presence was also confirmed by the observation of GFP-positive structures in the kidney parenchyma and a 3-fold GFP copies concentration compared to the control, perfused kidney. Our results are in line with that published in 2020 by Thompson et al^41^. In addition, at 6 hours, the treated kidney graft showed a lower expression of kidney injury and inflammatory markers compared to the control, as well as a slightly higher urine output. These findings suggest that infused tubuloids may contribute to reducing ischemia-reperfusion injury.

Our study has some limitations. Although it suggests that tubuloids can be efficiently delivered and integrated into a kidney host, we haven’t assessed if they have a positive impact on kidney function since the transplantation model was performed to assess biodistribution and engraftment, and no native nephrectomy was performed in the rat recipient. In the human kidney analysis, although we observed a potential positive effect of tubuloid infusion on expression parameters, only one case was performed, so solid conclusions cannot be precluded. Thus, further studies to assess the potential benefits of tubuloid implantation in kidney conditioning and improvement are needed.

In conclusion, our work reinforces the role of tubuloids as a key point for studying kidney diseases and developing new treatments, as well as promoting kidney tubular regeneration. Moreover, it provides evidence about the engraftment ability of tubuloids and raises NMP as an efficient technique for regenerative cell therapy that overcomes the off-target delivery to other organs. Since NMP is a platform already developed for solid human organs in clinical practice, this strategy may become a promising approach for kidney conditioning and regeneration with fast translatability to clinical practice. Future research in regenerative strategies is necessary to exponentially expand the knowledge in tissue regeneration and rapidly translate these therapeutic advances into clinical practice.

## ACKNOWLEDGEMENTS

This study has been founded by Klerk Investigations AIE, Instituto de Salud Carlos III through the Rio Hortega Contracts (project CM22/00172, funded by Instituto de Salud Carlos III and co-funded by the European Union NextGenerationEU/Mecanismo para la Recuperación y la REsilencia (MRR)/PRTR), the project “PI21/00205” (funded by Instituto de Salud Carlos III and co-funded by the European Union), the project RD21/0005/0003 (funded by Instituto de Salud Carlos III and co-funded by the European Union NextGenerationEU/Mecanismo para la Recuperación y la REsilencia (MRR)/PRTR) and by the Emili Letang-Josep Font grant from the Hospital Clinic of Barcelona.

## DISCLOSURE

The authors of this manuscript have no conflicts of interest to disclose.

## DATA AVAILABILITY STATEMENT

Data was generated in the Experimental Laboratory of Nephrology and Transplantation (IDIBAPS). Transcriptomic and imaging data supporting the findings of this study are included within the manuscript and the provided Supporting Information. All data generated or analyzed during this study are available from a Github repository. Datasets analyzed during the current study are available in the repository Sequence Read Archive (SRA) in the following Bioproject of NCBI PRJNA1153012.

## SUPPORTING INFORMATION STATEMENT

Additional supporting information may be found online in the Supporting Information section.

## Notes

### Competing Interest Statement

The authors have declared no competing interest.

