## Supporting Information for "Enhanced Kidney Targeting and Distribution of Tubuloids During Normothermic Perfusion"

**SUPPLEMENTAL METHODS**

Human tubuloid culture

Kidney tissue was mechanically minced and digested with a solution of DMEM-F12 (Gibco^TM^) containing 1.5 mg/mL collagenase (from Clostridium histolyticum, Sigma-Aldrich) and Bovine Serum Albumin (Sigma-Aldrich). The digested tissue was sequentially sieved through a 320 µm and 150 µm sieves. After a brief centrifugation, the kidney extract was resuspended in Matrigel^®^ (Corning) and plated to obtain tubuloids (protocol A, Crude tubuloids). We performed a second protocol (protocol B, F4 tubuloids) in which, after completing protocol A, the tissue further underwent a second separation process by centrifugation with 45% Percoll (Sigma-Aldrich) to create a density gradient resulting in four distinct bands (F1 – F4) as described previously^17^. F1 – F3 were discarded and only F4 (an enriched fraction with proximal tubular cells) was collected, resuspended and cultured in Matrigel^®^ to obtain tubuloids.

After including this purification step, tubuloid culture and development (for both F4 and Crude tubuloids) were performed as previously described^7^. Briefly, tubular cells were cultured in a tridimensional way using growth factor-reduced Matrigel® (Corning). Culture medium was composed of Advanced DMEM (Gibco^TM^) supplemented with 1% penicillin/streptomycin (Gibco^TM^), 1% HEPES (Gibco^TM^), 1% GlutaMAX (Gibco^TM^), N-acetylcysteine (1 mM, Sigma), 1.6% B27 supplement (Gibco^TM^), 1% R-Spondin 3 conditioned medium (U-protein express BV), human EGF (50 ng/ml, Peprotech), and human FGF-10 (100 ng/ml, Preprotech), Rho-kinase inhibitor Y-27632 (10 μM, Abmole), A8301 (5 μM, Tocris Bioscience) and primocine (0.1 mg/ml, Invivogen). The medium was changed every other day, and tubuloids were expanded in a 1:4 ratio once per week. For expansion, tubuloids were mechanically disrupted with a 27G needle before seeding.

Rat tubuloid culture

Rat kidney tissue for rat tubuloid culture was collected from healthy 3 months-old Lewis rats (LEW/Han®Hsd, Envigo^®^). The animals were kept at constant temperature, humidity, and at a 12-hour light/dark cycle with free access to water and rat chow until sacrifice. Kidney tissue was obtained after nephrectomy of both kidneys: a double nephrectomy was performed under general anesthesia and the kidneys were immediately placed in Celsior^®^ at 4ºC until processing. Rats were immediately euthanized under anesthesia. The study was approved by and conducted according to the guidelines of the local animal ethics committee (Comité Ètic d’Experimentació Animal, CEEA, Decret 214/97, Catalonia, Spain).

Tubular cell enrichment before culture was performed as described for human tubuloids: from the same rat donor, one kidney was processed through the Crude protocol and the contralateral through the F4 protocol. Tubuloid culture and development were performed following the same methods that those described for human tubuloids, although rat FGF (100 ng/ml, Preprotech) and rat EGF (50 ng/ml, Peprotech) were used instead of human FGF and EGF.

Tubuloid infusion during rat kidney normothermic perfusion

Rat donor organ procurement was performed under anesthesia with isoflurane (IsoVet®, Braun) as previously described^18^. Briefly, donor kidneys were flushed with sterile physiologic saline at 4°C after systemic application of sodium heparin (1000 UI, Rovi) and then the nephrectomy was performed. Renal artery and vein were carefully isolated, and other small vessels were ligated to avoid perfusate leaking during perfusion.

For rat kidney normothermic perfusion, Hugo Sachs/Harvard Apparatus device was used following the protocol published elsewhere^19^. Briefly, this system consists of a water-jacketed and circular chamber with a glass lid assembly. The kidney is connected through the cava vein and the aorta using two cannulae for organ perfusion. Associate with this main part, the system includes different support components to maintain the organ in the most physiological conditions: a membrane oxygenator, reservoirs, cannula line holders, perfusate, gas and water lines, bubble traps and manifolds for water, gas and perfusate control. The perfusate is pre-warmed at 37ºC and oxygenated for 30 minutes before organ connection. Perfusate was composed of Gey’s solution (Sigma) and 1% Penicillin/Streptomycin (Sigma). After organ connection, a pressure-controlled perfusion was performed, increasing progressively the pressure for 5 minutes to reach a mean pressure of 70 mmHg. Nevertheless, for those kidneys in which a high vascular resistance was observed, a minimum flow of 0.5 mL/min was set.

GFP-expressing rat tubuloids were harvested and resuspended in pure Matrigel® with previous mechanical disaggregation and observed under the microscope to ensure a 25-50 µM diameter before infusion. After stabilizing the kidney graft, tubuloids were administered in the perfusate through the arterial line. Kidney grafts were maintained under NMP for 1 hour. After finishing NMP, epifluorescence was immediately assessed with an IVIS® Lumina III (Perkin Elmer) and then processed for PCR and immunofluorescence analysis.

For each donor, one kidney graft was perfused and treated with tubuloids, while the contralateral one was perfused but no tubuloids were administered. We tested two doses of tubuloids: 1·10^6^ cels/g (high dose, n = 4 kidneys) and 3·10^5^ cels/g (low dose, n = 3 kidneys).

Kidney Transplantation in Rats

Briefly, an end-to-side anastomosis of the aortic stump of the donor kidney and recipient’s aorta, and between the recipient inferior vena cava and donor renal vein, respectively, using 9-0 running nylon sutures. The mean time for anastomosis was 24.8 ± 3.3 minutes. Uretero-ureterostomy was performed with an end-to-end interrupted sutures technique using 11-0 nylon sutures. Native kidneys were not removed at the time of engraftment. 1 week and 1 month after transplantation, recipient rats were euthanized and native kidneys, kidney graft and liver were retrieved for analysis.

Tubuloid infusion during human kidney normothermic perfusion

The renal artery was cannulated to the circuit using a 12-Fr cannula; renal vein outflow was via pump to an oxygenator whereby blood re-entered the main circuit. The ureter was directly cannulated to the urine circuit and collected without recirculation. A urine flow sensor monitored urine output. Instead of urine recirculation, repositioning with a balanced solution (Isofundin®) according to urine output was conducted with a pump connected to the urine flow sensor. Hemodynamics were pressure-controlled, establishing a pressure increase ramp until reaching a set mean arterial pressure of 70mmHg during perfusion. The temperature was set at 37ºC during the perfusion.

The system was primed with two units of packed isogrup Red Blood Cells (RBC) and 500mL of balanced saline solution (Isofundin®), which an approximate total volume of 800mL. Oxygenation of the perfusate was performed by manual regulation of air (21% oxygen) at a flow rate of 1.5mL/min to maintain an oxygen saturation over 97% using an ECMO oxygenator. A nutrition solution was continuously infused at 24 mL/h. Nutrition was composed of 10% Aminoplasmal®, Cernevit® as a multivitamin complex, 5% Serum glucose and 70UI of insulin. Verapamil was used as a vasodilator (continuous infusion rate, 0.4 mg/h). Before kidney connection to the circuit, 1.2g of amoxicillin-clavulanate, 10 mL of sodium bicarbonate 8.4%, 8mg of dexamethasone, 2000UI of low molecular weight heparin, 5 mL of glucose 5% and 0.5 mL of calcium gluconate were added. The same dose of antibiotic was administered every 24 hours. 10mL of sodium bicarbonate 8.4% was administered to maintain a pH >7.30. Finally, 5mL of glucose 5% was administered as needed to maintain a perfusate glucose >100 mg/dL.

Biochemical parameters were measured in the perfusate using the Epoc® Blood Analysis System (Siemens Healthcare). Perfusate samples were collected hourly. Histological samples were collected at the beginning of the perfusion (time point 0) and at the end of the perfusion (6 hours). A 1x1x1mm piece of kidney tissue was taken from the middle third of the graft and immediately processed. In case of bleeding after the biopsy, the tissue was carefully sutured to avoid leakage and ensure that renal flow was not affected.

Both kidneys were maintained for 6 hours. Once NMP was finished, GFP fluorescence was immediately assessed in an IVIS® Lumina III (Perkin Elmer) and then processed for PCR and immunofluorescence analysis.

Histology and immunofluorescence analysis

Tubuloids were fixed in 4% paraformaldehyde (ThermoFisher Scientific), included in 8% agarose (UltraPure^TM^ Low Melting Point Agarose, ThermoFisher Scientific) and then embedded in paraffin. Afterwards, 3 µm sections were stained with Hematoxylin-Eosin (H&E) or different antibodies for immunofluorescence assays.

The following antibodies were used: Rabbit anti-AQP-3 (1:250, 125219, Abcam), mouse anti-Calbindin (1:250, 365360, Santa Cruz), rabbit anti-E-Cadherin (1:250, 610182, BD Biosciences), rabbit anti-Integrin ɑ6 (1:250, AB20142, Abcam), mouse anti-Villin (1:250, SC-58897, Santa Cruz), rabbit anti-ZO1 (1:250, 402200, Thermo Fisher Scientific), mouse anti-acetylated tubulin (1: 250, SC-23950, Santa Cruz), rabbit anti-cytokeratin (1:500, ab9377, Abcam) and rabbit anti-Aquaporin 3 antibody (Anti-AQP3, AB125219, Abcam) were used as primary antibodies. Alexa Fluor immunoglobulin Gs (IgGs) were used as secondary antibodies: 488 donkey anti-rabbit IgG (1:500, 711545152, Jackson Immunoresearch) and 555 donkey anti-mouse IgG (1:500, A-31570, Thermo Fisher Scientific). Nuclei were stained with immunomounting media with DAPI (Thermo Fisher Scientific). A primary anti-PAX8 combined with a fluorochrome was used for PAX-8 immunofluorescence (1:100, ab217733, Abcam). Images were obtained with a spectral high-speed confocal microscope (Zeiss LSM880) and processed using Image J (U. S. National Institutes of Health, Bethesda, Maryland, USA). Histology images were acquired using a Leica DMI6000 B inverted microscope (Leica Microsystems, Germany).

Organ tissues were fixed in formaldehyde (Sigma) for 24 hours and then embedded in paraffin. Afterward, 3-µm sections were performed. To assess GFP tubuloid implantation in the kidney host, an immunofluorescence GFP analysis using a GFP nanobody (ChromoTek GFP-Booster ATTO488) was performed according to the manufacturer’s protocol. Nuclei were stained with immunomounting media with DAPI (Thermo Fisher Scientific). A protocol to mitigate kidney autofluorescence was performed as previously described^21^. Immunofluorescence images were acquired using an Olympus BX51 Fluorescence Microscope (Leica Microsystems) under the same acquisition and detection settings. GFP signal was corrected for background.

Quantitative Polymerase Chain Reaction (qPCR)

Tubuloid samples were lysed and homogenized and total RNA was extracted using Maxwell® RSC Instrument (Promega) with the Maxwell® RSC miRNA from Plasma or Serum kit (Promega). For tissue RNA extraction, the Maxwell® RSC miRNA from Tissue (Promega) was used, according to the supplier’s protocol.

cDNA was synthesized from the RNA template using a High Capacity cDNA Reverse Transcription Kit (Applied Biosystems™ 4368813) as per the manufacturer’s instructions. The resulting cDNA was used to determine expression levels.

Real-Time qPCR was performed using the corresponding TaqMan® gene Expression Assay (20x) (Invitrogen, Supplemental Table S2) and TaqMan Fast Advanced Master Mix (Applied Biosystems) on a 384-well plate using the PCR program provided by the supplier, on a QuantStudio 7 device (ThermoFisher). Samples were run in triplicate in 10 μl reactions, and mRNA expression of the target genes was normalized to Beta-Actin mRNA and expressed as the fold change with the ΔΔCT method, according to the model. Expression levels of the following genes were measured: ABCC4, SLC12A1, SLC12A3, SLC41A3 and AQP3.

For human kidney tissue, real-time qPCR was performed using the corresponding primers for each gene on a 384-well plate using the PCR program provided by the supplier. Samples were run in triplicate in 10 μl reactions, and mRNA expression of the target genes was normalized to β-actin mRNA and expressed as the fold change to time 0 kidney tissue with the ΔΔCT method. Expression levels of KIM-1, CASP3. TNFa and VEGF were measured (Supplemental Table S3).

Digital PCR (dPCR)

The dPCR was set up according to manufacturer instructions (Applied Biosystems). The Applied Biosystems QuantStudio Absolute Q Digital PCR System was used. This system is a plate-based dPCR platform with unique microfluidic array plate technology that enables all steps required for dPCR (compartmentalization, thermal cycling and data acquisition) in a single instrument. All reactions were performed and read on a 20-well plate. The condition of dPCR was optimized as per manufacturer recommendations: 95 °C for 10 min, 40 × (96°C for 5s, 60°C for 15 s). A TaqMan EGFP probe (Invitrogen, Supplemental Table S1) was used for copy quantification in cDNA samples obtained as described before. Absolute Q™ DNA Digital PCR Master Mix (5X, Invitrogen) was used according to the manufacturer’s protocol. Data was read and analyzed using the QuantStudio Absolute Q Digital PCR software.

Single-cell RNA sequencing (scRNA)

scRNA sequencing was performed with tubuloids obtained from 3 different patients. Passage 3 human tubuloids were grown for 1 week post-dissociation. After that, disaggregation and Matrigel® removal were performed by treatment with 1X TrypLE Express (Gibco) for 5 min at 37ºC. Tubuloids were further dissociated mechanically using a 27G needle. When single cells were obtained, tubuloids were washed, counted and resuspended in PBS with non-acetylated BSA (UltraPure™ BSA Non-Acetylated, ThermoFisher).

Single-cell RNA sequencing data processing, filtering, and clustering

Single-cell RNA sequencing was performed using the 10x Genomics Chromium platform. Sample reads were aligned to the human genome GRCh38 using the Cell Ranger software suite (10x Genomics). This process included the steps of demultiplexing, aligning reads to the reference genome, and quantifying gene expression levels by generating UMI (Unique Molecular Identifier) counts. The result was a gene expression matrix that served as the basis for subsequent analysis.

All downstream analyses were conducted using the Seurat package in R. Initially, the gene expression matrix produced by Cell Ranger was imported into R and converted into a Seurat object. Cells were filtered based on quality control metrics to remove low-quality cells and potential multiplets. Specifically, cells with fewer than 200 or more than 7500 detected transcripts were excluded, as were cells with more than 15% of reads mapping to mitochondrial genes. This stringent filtering ensured the retention of high-quality single-cell data.

Filtered datasets from multiple samples were then integrated using reciprocal principal component analysis (RPCA) to mitigate batch effects and harmonize the data. Integration anchors were identified across datasets, and data were combined into a single Seurat object. The integrated data were then log-normalized to stabilize variance across genes.

Dimensionality reduction was performed using principal component analysis (PCA). The top 20 principal components, which explained the majority of variance in the data, were selected for downstream analysis. Clustering was subsequently performed using the FindNeighbors function, which constructs a shared nearest neighbour graph based on the PCA-reduced data (dimensions 1:20), followed by the FindClusters function, which applies the Louvain algorithm for community detection at a resolution of 0.5. This method enabled the identification of distinct cell clusters within the dataset.

To visualize the clustering results, Uniform Manifold Approximation and Projection (UMAP) was employed, providing a two-dimensional representation of the high-dimensional data. Clusters were annotated by identifying differentially expressed genes using the FindAllMarkers function, applying a log fold-change threshold of 0.25 to highlight genes that were significantly upregulated in each cluster.

In the analysis of Crude tubuloids, an elbow plot was used to determine the optimal number of clusters. The elbow plot, which plots the total within-cluster sum of squares against the number of clusters, indicated that two clusters were optimal for the dataset. This method helps in identifying the point where the addition of more clusters does not significantly reduce the within-cluster variation, signifying a natural grouping in the data. Consequently, the analysis resulted in the identification of two distinct clusters in both Crude and F4 tubuloids

For the combined gene expression analysis and Z-scoring of Crude and F4 clusters, data were obtained in CSV format, containing measurements across the two experimental conditions (Crude and F4) and two biological replicates for each condition (Crude 0, Crude 1, F4 0 and F4 1). The dataset also included a column representing gene identifiers (Gene). For downstream analysis, only these columns were selected, and the Gene column was set as the index to streamline data handling.

To standardize gene expression values across the experimental conditions, Z-scores were computed for each gene individually. Z-score normalization was performed by subtracting the mean expression value of each gene and dividing by its standard deviation. This approach ensured that the data were centered and scaled, facilitating a comparative analysis of relative gene expression levels across conditions. The normalized Z-scores were visualized using a heatmap to highlight patterns of gene expression regulation across conditions.

**SUPPORTING FIGURES**

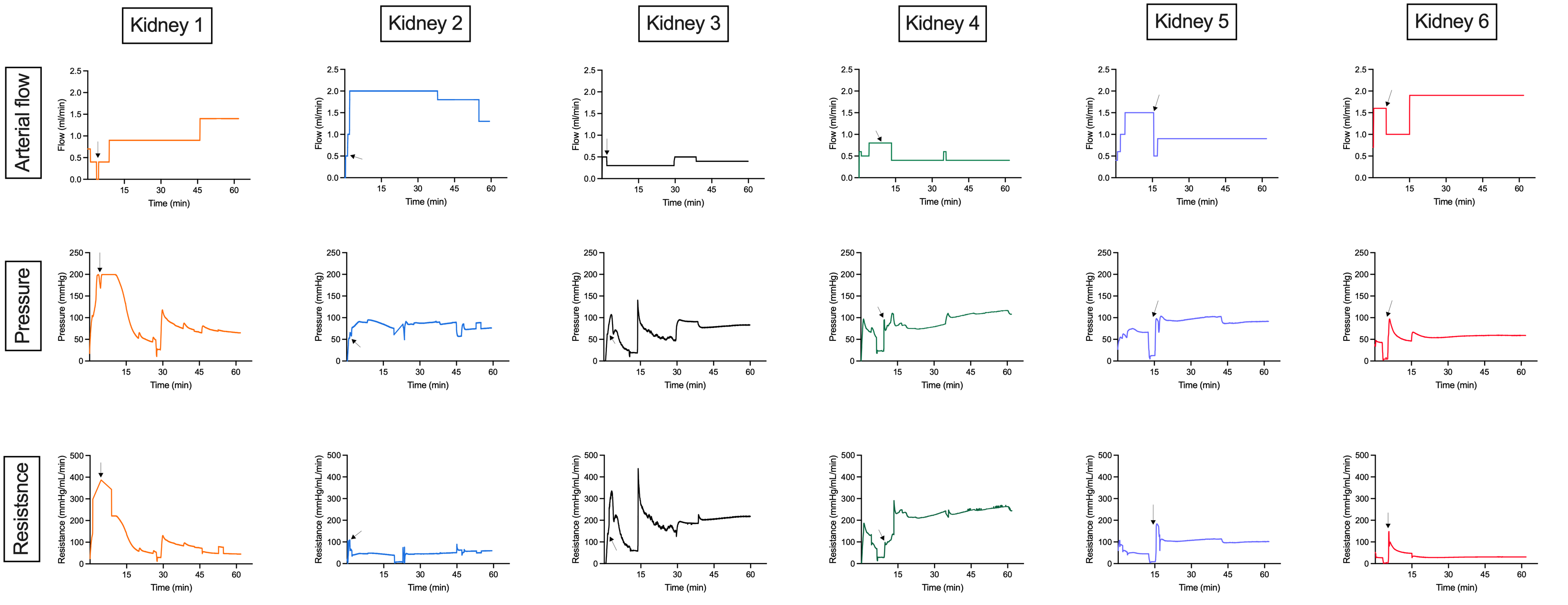

**Supplementary Figure S1. Rat Kidney hemodynamics during normothermic perfusion.** Arterial flow, pressure and resistance during normothermic perfusion for 6 representative cases. Kidneys 1 to 4 were only used for immediate biodistribution assessment, without transplantation. Kidney 5 and 6 were transplanted in a recipient rat. Black arrows indicate the moment in which tubuloids were administered to the perfusate.

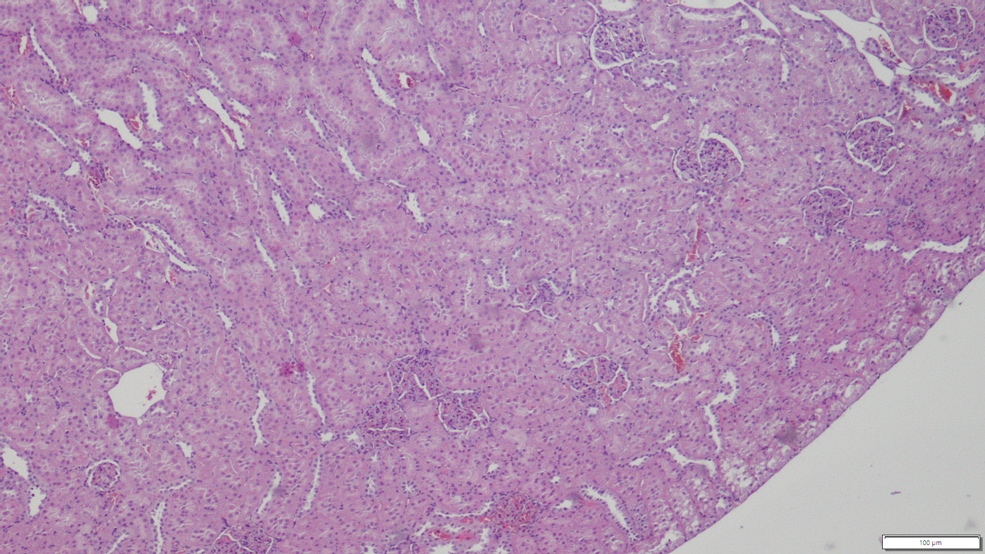

B

A

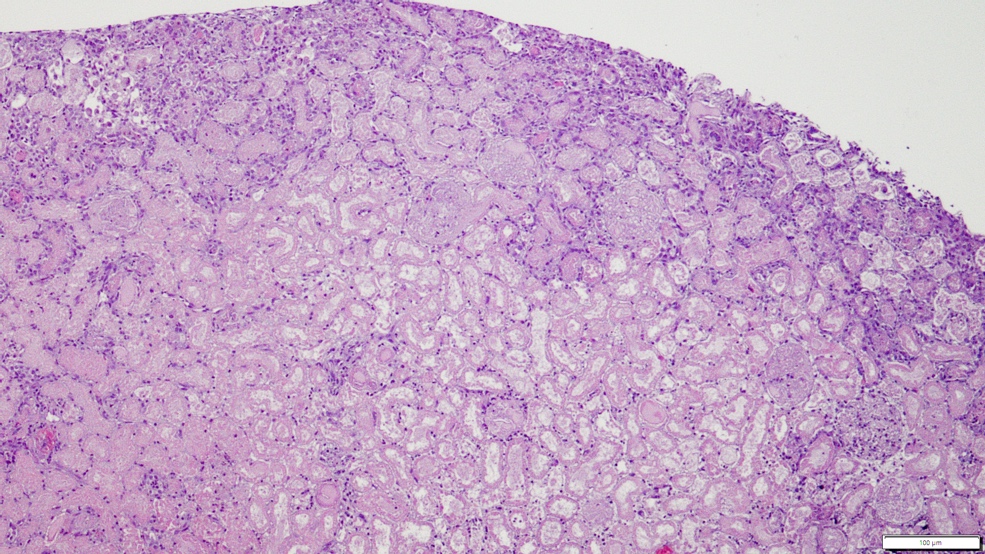

C

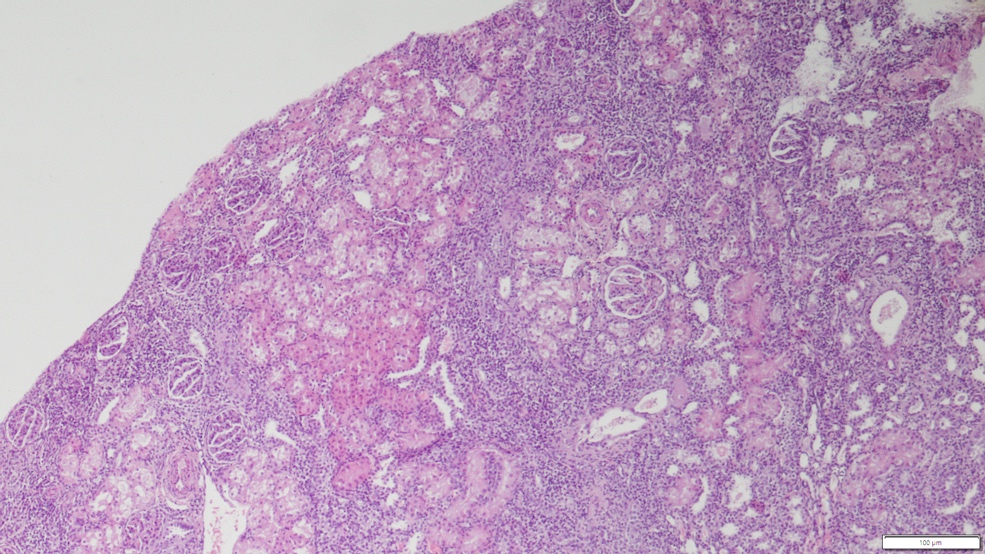

**Supplementary Figure S2. Hematoxylin-Eosin Staining of native and transplanted kidneys during normothermic perfusion after 1 week and 1 month.** A, Contralateral kidney (no treated, no perfused)(n = 4 rat recipients). B, Kidney graft treated with tubuloids during normothermic perfusion and collected 1 month after transplantation (n = 4 rat recipients). C, Kidney graft treated with tubuloids during normothermic perfusion and collected 1 month after transplantation (n = 4 rat recipients).

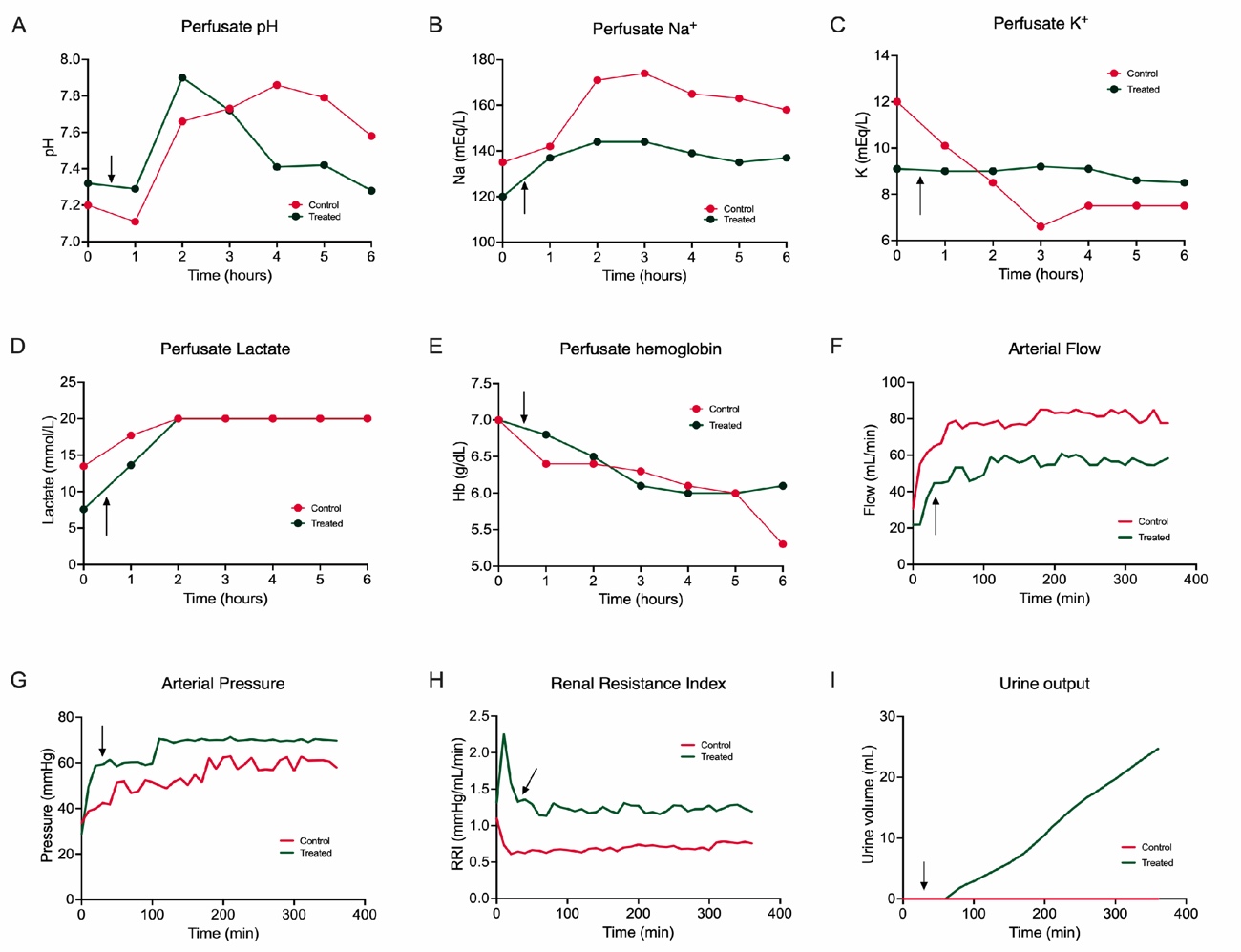

**Supplementary Figure S3. Human Kidney hemodynamics and biochemistry during normothermic perfusion.** A, Perfusate pH; B, Perfusate Na; C, Perfusate K; D, Perfusate lactate; E, Perfusate hemoglobin; F, Arterial flow during normothermic perfusion; G, Arterial pressure during normothermic perfusion; H, Vascular resistance during normothermic perfusion; I, Urine output during normothermic perfusion. N = 1 kidney per group (control and treated).

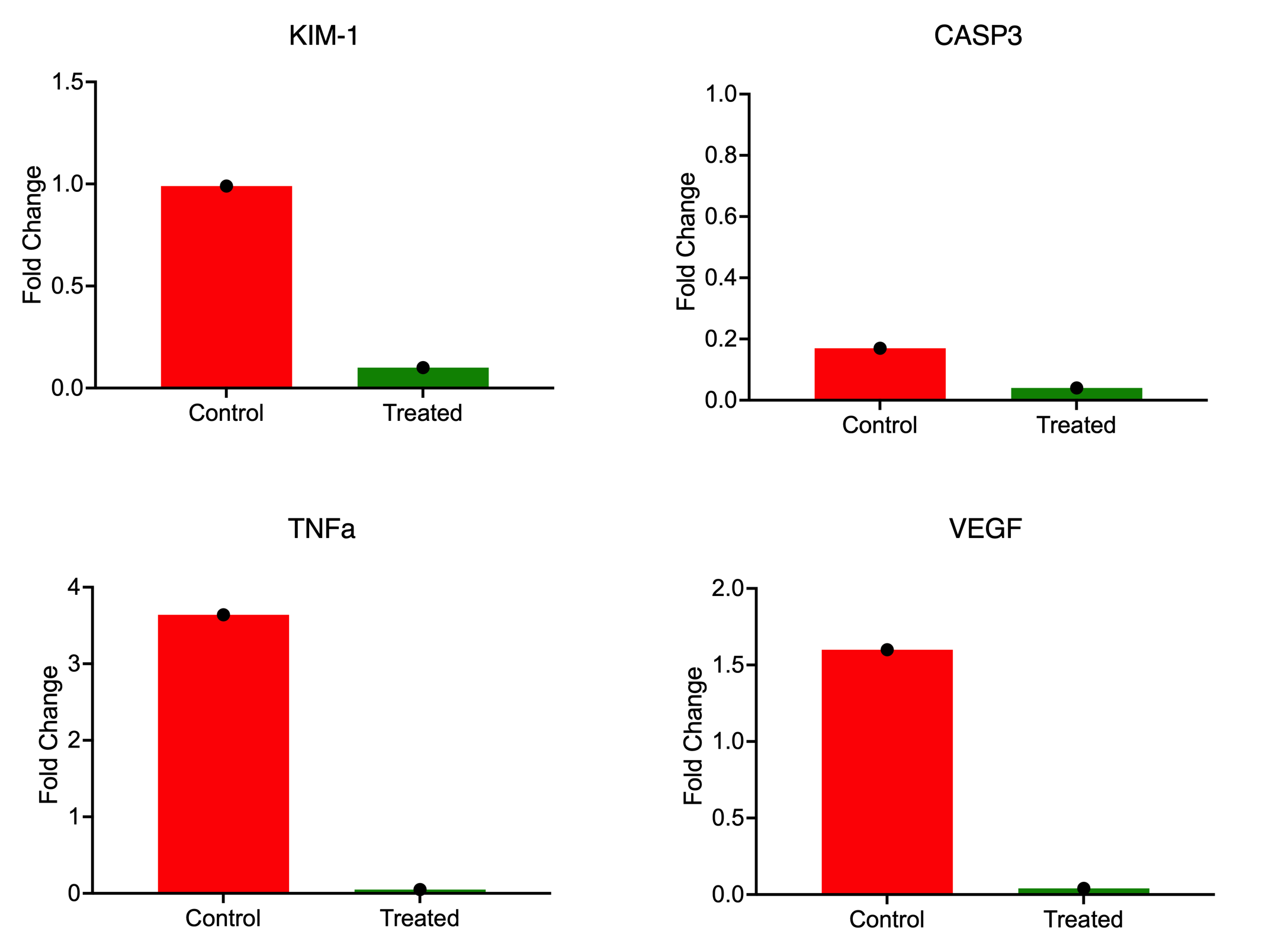

**Supplementary Figure S4. Human Kidney expression of kidney injury and inflammatory markers during normothermic perfusion in treated and control human kidneys.** Gene expression has been represented as fold change at 6 hours of perfusion for time 0. KIM-1, Kidney Injury Marker 1; CASP3, Caspase 3; TNFa, Tumor Necrosis Factor alfa; VEGF, Vascular Endothelial Growth Factor. N = 1 kidney per group (control and treated). Each sample was run in triplicates.

**SUPPORTING TABLES**

| **CRUDE TUBULOIDS** | |
| --- | --- |
| **Gene** | **Location** |
| RPA1 | Collecting duct |
| FEM1A | Proximal Tubule |
| SDK1 | Proximal Tubule |
| NALF1 | Collecting duct |
| NHS | Proximal Tubule |
| CACNA2D3 | Distal Tubule |
| FHIT | Proximal Tubule |
| PTPRM | Proximal Tubule |
| KAZN | Collecting duct (mostly) |
| GLIS3 | Proximal and collecting duct (equal) |
| IMMP2L | Distal and proximal tubules |
| JMJD1C | Proximal tubule and collecting duct |
| **F4 TUBULOIDS** |  |
| S100A6 | Thick Ascending Limb (progenitors) |
| S100A11 | Proximal/Distal/Collecting duct |
| RPL41 | Proximal/Distal/Collecting duct |
| SPP1 | Proximal |
| EEF1A1 | Proximal |
| FTH1 | Proximal |
| TPT1 | Proximal |
| RPLP1 | Proximal (mostly) |
| TMSB4X | Non-defined |
| RPS12 | Proximal and distal tubules |
| NALF1 | Collecting duct |
| ERBB4 | Distal Tubule and Collecting duct |
| PTPRM | Proximal |
| NEAT1 | Distal |
| MALAT1 | Non-defined |
| KCNIP4 | Distal Tubule |
| MECOM | Distal Tubule |
| LINC00472 | Non-defined |
| EFNA5 | Distal Tubule |
| BICC1 | Proximal and Distal Tubules |

**Table S1**. Genes analyzed in scRNA analysis for Crude and F4 tubuloids (data extracted from Human Protein Atlas proteinatlas.org and Karlsson M et al Sci Adv. 2021 Jul)

| **Gene Symbol** | **Gene Name** | **Detector** | **Location** |
| --- | --- | --- | --- |
| Human ABCC4 | ATP Binding Cassette Subfamily C Member 4 | Hs00988721_m1 | Proximal Tubule |
| Human SLC12A1 | Solute Carrier Family 12 member 1 | Hs00165731_m1 | Ascending limb Henle Loop |
| Human SLC12A3 | Solute Carrier Family 12 member 3 | Hs01027568_m1 | Distal Tubule |
| Human SLC41A3 | Solute Carrier Family 41 Member 3 | Hs00215208_m1 | Distal Tubule |
| Human AQP3 | Aquaporine 3 | Hs00185020_m1 | Collecting Duct |
| Rat ABCC4 | ATP Binding Cassette Subfamily C Member 4 | Rn01465702_m1 | Proximal Tubule |
| Rat SLC12A1 | Solute Carrier Family 12 member 1 | Rn00692576_m1 | Ascending limb Henle Loop |
| Rat SLC12A3 | Solute Carrier Family 12 member 3 | Rn01531762_m1 | Distal Tubule |
| Rat SLC41A3 | Solute Carrier Family 41 Member 3 | Rn01426072_m1 | Distal Tubule |
| Rat AQP3 | Aquaporine 3 | Rn00581754_m1 | Collecting Duct |
| EGFP | Enhanced GFP | Mr04329676_mr | N/A |

**Table S2**. Gene list of Taqman Expression Assays

| **Gene** | **Forward** | **Reverse** |
| --- | --- | --- |
| Human KIM-1 | 5’-TGGCAGATTCTGTAGCTGGTT-3’ | 5’-AGAGAACATGAGCCTCTATTCCA-3’ |
| Human CASP3 | 5’-TTTGAGCCTGAGCAGAGACA-3’ | 5’-CGTATGGAGAAATGGGCTGT-3’ |
| Human TNFa | 5’-ACTTTGGAGTGATCGGCC-3’ | 5’-GCTTGAGGGTTTGCTACAAC-3’ |
| Human VEGF | 5’-GAATGGGGAGCCCAGAGTG-3’ | 5’-TCCTCCTTCTGCCATGGGT-3’ |
| Human B-actin | 5’-CCTCGCCTTTGCCGATCC-3’ | 5’-CGCGGCGATATCATCATCC-3’ |

**Table S3**. List of primer sequences for qPCR Assays
